# *Cx3cr1*-BAC-CRE-mediated knockout of toll-like receptor 4 alters mouse communicative behaviors, microglial morphology, and engulfment of synaptic material

**DOI:** 10.64898/2026.09.17.752150

**Authors:** Izabella M Bankowski, Indra R Bishnoi, Eva E Shin, Mia J Cashin, Dorit Möhrle, Haley A Norris, Drew T Barry, Maggie M Slamin, Evan A Bordt

## Abstract

Microglia, the tissue resident immune cells of the central nervous system, are critical regulators of postnatal neural circuit refinement. Innate immune receptors on microglia such as toll-like receptor 4 (TLR4) are best characterized in the context of inflammation, however, endogenous TLR4 ligands are generated during typical developmental processes. Here, we investigated whether TLR4 signaling in CX3CR1-expressing myeloid cells contributes to microglial morphology, synaptic engulfment, and behavioral development in mice under basal conditions. TLR4 conditional knockout altered maternal separation-induced ultrasonic vocalizations without impacting social preference or anxiety-like behaviors. Conditional TLR4 knockout in CX3CR1-expressing myeloid cells markedly increased microglial ramification and cell volume within the paraventricular nucleus (PVN) of the hypothalamus. TLR4 conditional knockout also reduced microglial engulfment of vGlut2-positive presynaptic material, while overall excitatory synapse numbers remained unchanged. Together, these findings demonstrate that TLR4 signaling in CX3CR1-expressing myeloid cells contributes to microglial morphology, presynaptic engulfment, and neonatal communicative behavior under basal conditions.

## INTRODUCTION

Microglia, the tissue resident immune cells of the central nervous system, refine neural circuits across development through activity-dependent synaptic pruning. During postnatal development, microglia selectively engulf synaptic inputs in a complement-dependent manner (Stevens et al., 2007; Schafer et al., 2012). Dysregulation of synaptic pruning during sensitive developmental windows has emerged as a potential mechanism linking early-life adversity to enduring dysfunction in stress-related neural circuitry. Toll-like receptor 4 (TLR4) is a pattern recognition receptor that is canonically appreciated for its role in detecting exogenous pathogen-associated molecular patterns (PAMPs) such as lipopolysaccharide (LPS) and initiating innate immune signaling cascades. In addition to PAMPs, TLR4 can also recognize a diverse array of endogenous ligands and damage-associated molecular patterns (DAMPs) including HMGB1 (Guazzi et al., 2003; Zhao et al., 2011, 2020), S100 proteins (Masuda et al., 1983; Tramontina et al., 2002; Hachem et al., 2007), and heat-shock proteins (Luft and Dix, 1999; David et al., 2001). Within the developing brain, these endogenous ligands are generated through physiological processes such as apoptosis, synaptic refinement, and extracellular matrix turnover, positioning TLR4 as a sensor of developmental cues in addition to its role as a sensor of infection or injury. Since developmental tissue remodeling continuously generates these TLR4 ligands, tonic TLR4 signaling may provide a mechanism by which microglia monitor and respond to normal circuit remodeling throughout postnatal development. Indeed, developmental TLR4 deficiency has been demonstrated to enhance cognitive effects correlated with CREB upregulation in the hippocampus (Okun et al., 2012), suggesting that TLR4 may influence neural circuit organization even in the absence of overt immune challenges. Furthermore, TLR4 expression on microglia is required for normal phagocytic responses to endogenous extracellular matrix components expressed during development (Song and Dityatev, 2018; Haage et al., 2019; Yang et al., 2023), highlighting a key role for TLR4 in regulating homeostatic function during brain development.

Beyond regulating inflammatory responses, TLR4 also influences fundamental aspects of microglial biology. Microglial morphology and surveillance are actively maintained by receptor-dependent signaling (Davalos et al., 2005; Nimmerjahn et al., 2005; Madry et al., 2018). Accordingly, TLR4 activation alters microglial ramification (Papageorgiou et al., 2016), suggesting that TLR4 contributes to the regulation of microglial structural and homeostatic functions. However, whether TLR4 regulates these developmental functions under physiological conditions remains poorly understood.

Despite these converging lines of evidence, the field has predominantly studied TLR4, and indeed microglial innate immune receptors more broadly, within the framework of pathological activation. Most experimental paradigms examine TLR4 function under conditions of exogenous immune challenge, neurological injury, disease, or stress. Therefore, the extent to which TLR4 signaling shapes microglial morphology, synaptic activity, and downstream behavioral outcomes under basal conditions remains poorly understood. As microglia actively sculpt developing neural circuits, perturbations of microglial function during sensitive developmental windows alter not only synaptic connectivity but also behavioral outcomes that reflect the integrity of these circuits (Branchi et al., 2001; Scattoni et al., 2009). Notably, microglial dysfunction during the perinatal period has been shown to alter both synaptic connectivity and social communicative behaviors (Zhan et al., 2014; Kopec et al., 2018). Whether TLR4 within microglia is required for the synaptic engulfment that characterizes normal circuit refinement, and whether microglial TLR4 contributes to behavioral development under basal conditions has, to our knowledge, not been examined.

CX3 motif chemokine receptor 1 (CX3CR1, also known as GPR13) is highly expressed by microglia and mediates neuron–microglia interactions through its ligand CX3CL1 (fractalkine) (Jung et al., 2000; Paolicelli et al., 2014; Pagani et al., 2015). Cre-LoxP-mediated targeting of CX3CR1-expressing myeloid cells is a widely used method in microglial biology. To address these questions, we generated mice with conditional knockout of TLR4 in CX3CR1-expressing myeloid cells (TLR4 cKO) and assessed microglial morphology, synaptic engulfment in the paraventricular nucleus (PVN) of the hypothalamus due to its established role in stress-sensitive microglial pruning (Bolton et al., 2022; Garvin et al., 2025). Since neonatal ultrasonic vocalizations (USVs) provide an early-life readout of neural circuit function during the same developmental window during which microglia undergo synaptic remodeling, we examined maternal separation-induced USVs as an early behavioral consequence of altered microglial TLR4 signaling. To assess whether the consequences of altered TLR4 signaling persist into later behavioral domains, and since developmental alterations in microglial function have also been linked to lasting changes in social and anxiety-like behaviors (Zhan et al., 2014; Nelson and Lenz, 2017; Jones et al., 2025), we also assessed adolescent social and anxiety-like behaviors. We demonstrated that TLR4 cKO altered communicative behaviors, produced marked alterations in microglial morphology, and significantly disrupted the engulfment of excitatory synaptic components by microglia in the PVN. These findings demonstrate that TLR4 in CX3CR1-expressing myeloid cells is required for normal microglial morphology, presynaptic engulfment, and neonatal communicative behavior under basal conditions, supporting a physiological role for innate immune signaling during postnatal circuit refinement.

## METHODS

### EXPERIMENTAL MODEL AND SUBJECT DETAILS

#### Experimental Animals

TLR4^flox/flox^ mice were bred with *Cx3cr1*-BAC-Cre mice (Ceasrine et al., 2022; Bordt et al., 2024) to generate conditional knockout of TLR4 in CX3CR1-expressing myeloid cells (TLR4 cKO; Cre positive) and Cre negative fully floxed littermate controls. Mice were time-mated for 5 days using either TLR4^flox/flox^:Cre^+^ females x TLR4^flox/flox^:Cre^-^ males or TLR4^flox/flox^:Cre^-^ females x TLR4^flox/flox^:Cre^+^ males. Litters were mixed-genotype (Cre^+^ and Cre^-^). To confirm TLR4 knockout in microglia, microglial isolations were performed on forebrain tissue as previously described using CD11b microbead method (Bordt et al., 2020, 2024), after which TLR4 gene expression in CD11b+ or CD11b-cells was assessed by Taqman qPCR array.

Animals were bred in standard polypropylene cages (27 cm x 16 cm x 15.5 cm) with cobb bedding and CareFresh nesting material. Pregnancy checks were performed by visualization of vaginal plugs with an indication of a plug representing embryonic day 0.5 (E0.5). Upon identification of a plug, the dam was removed from the breeding pair and placed into an individual fresh polypropylene cages with cobb bedding, a cotton nestlet (1in x 1in square) in place of CareFresh, and an igloo for enrichment. Cages were subsequently exchanged for identical clean replacements on E14.5 (based upon vaginal plug date) and cages were inspected every morning beginning E17.5 to identify birth dates (postnatal day 0 – PND0).

Toe clips were performed on PND6 for animal identification, and tail snips were performed on PND18 for genotyping using PCR. Primers used for Tlr4 genotyping were: Forward: TGA CCA CCC ATA TTG CCT ATA C and Reverse: TGA TGG TGT GAG CAG GAG AG. After weaning, all mice were group housed in standard mouse cages under standard conditions (12-hour light/dark cycle, 23°C, 50% humidity) with same-sex littermates. Food pellets (Prolab IsoproRMH 3000 Irradiated Pellets) and tap water were available ad libitum, including during behavioral assays. All experiments were performed in accordance with the NIH Guide to the Care and Use of Laboratory Animals and approved by the Massachusetts General Hospital Institutional Animal Care and Use Committee.

### METHOD DETAILS

#### Maternal Observations

Maternal observations consist of noninvasive observations of maternal care behavior of dams after giving birth to identify altered patterns of maternal care delivery to pups. To evaluate whether the LBN paradigm was altering maternal care behaviors, we performed maternal observations of dams on PND3, PND6 and PND9. To do so, dams were observed starting from initial observation at 0 minutes (T0), and maternal behaviors were identified and catalogued every minute for the duration of 30 minutes. Whether or not dams were on the nest and performing typical nursing, licking or grooming behavior was noted.

#### Weaning

Mice were weaned individually by litter on PND24 into standard polypropylene cages (27 cm x 16 cm x 15.5 cm) with cobb bedding and CareFresh. Animals were housed only within matching sex and litter cages, with a maximum of 4 mice per sex per cage post-weaning based on IACUC guidelines.

### OFFSPRING NEONATAL BEHAVIORAL ASSAYS

#### Early Communicative Behavioral Evaluation via Ultrasonic Vocalizations

All behaviors are conducted across multiple cohorts due to the need for sex and genotype conditions. To evaluate early life communication, we recorded offspring maternal separation-induced ultrasonic vocalizations on PND8. We utilized a custom acrylic chamber padded on the inside with sound-proof foam on all four sides, with a hinged door. Inside the chamber, an ultrasonic microphone was mounted and secured to the center of the top of the chamber with wires exiting a hole on the roof of the chamber and connecting to an Avisoft Ultrasoundgate for recording (Ceasrine et al., 2022). PND8 pups were randomly placed into a plastic cup with 2 cotton pads. The plastic cup was cleaned with Peroxigard between each round. Two identical recording chambers were used simultaneously. All testing took place during the light phase. Mice were moved from their housing room to the behavioral room at approximately 8:00AM one hour prior to behavioral testing to habituate to the room. All recordings were automatically analyzed using VocalMat (Fonseca et al., 2021; Möhrle et al., 2023, 2024) and validated by a blinded scorer.

### OFFSPRING ADOLESCENT BEHAVIORAL ASSAYS

Adolescent behavioral assays were conducted at PND32-45 across multiple cohorts due to the need for sex and genotype conditions. All assays took place during the light phase and mice were moved from their housing room to the behavioral room one hour prior to behavioral testing to habituate to the room. Males and females underwent testing on separate days. Behavioral equipment was thoroughly cleaned with Peroxigard between trials.

#### Sociability Behavior

A 3-chamber social preference task was utilized to discern preference of experimental animals to investigate either a social stimulus (a novel age- and sex-matched conspecific) or a non-social stimulus (a rubber duck), also called sociability. For the first day of the task, animals were habituated to the chamber itself to eliminate it from being a novel environment for the experimental day. Animals were placed in the 3-chambered apparatus (60 cm total length divided in three 20 cm length chambers with a chamber width of 40 cm and wall height of 30 cm and an opening for movement between chambers) with empty round plexiglass containers (15 cm height, 10 cm diameter, 30 cm circumference, 22 vertical columns) on opposite end chambers of the apparatus and permitted to explore the apparatus for 5 minutes. Separately, novel age- and sex-matched conspecifics were habituated to the plexiglass cups for 5 minutes. On the subsequent test day, the sociability task was performed and recorded from both ceiling and side views. A novel age- and sex-matched conspecific and rubber duck were placed in round enclosed plexiglass containers and positioned in the opposite ends of the 3-chambered apparatus. Cups were aligned in the center openings into the chambers. Subsequently, the test animal was placed into the center chamber and allowed to explore for 5 minutes. Male and female behavior was run using two separate sex-specific 3-chamber test boxes. The following behavioral outcomes were scored manually by a blinded observer using Solomon Coder: time spent investigating the novel mouse and time spent investigating the novel object. Sociability was calculated as the amount of time spent investigating the novel mouse as a proportion of total time spent investigating both stimuli.

#### Anxiety-like Behavior: Elevated Zero Maze

As a test for the presence of anxiety-like behaviors, we utilized a zero maze (Stoelting) with elevated (50 cm) circulate lanes (5 cm wide) divided into four opposing sections. Two equally sized quadrants on opposite sides of the maze were enclosed by 15 cm tall walls on either side, identified as the “closed arm.” The other two evenly sized quadrants of the maze were 5 cm lanes without walls, identified as “open arms.” Test mice were placed into the center of the closed arm at the beginning of the test, and the time spent in the open and closed quadrants over the course of a 5-minute trial was recorded. Behavioral outcomes were scored manually by a blinded observer using Solomon Coder.

#### Anxiety-like Behavior: Light-Dark Box

As another measure of anxiety-like behavior, a light-dark box was utilized which consisted of a 35 cm tall box divided by a black plexiglass wall with an opening for movement between chambers. Each chamber was approximately 20 cm x 40 cm where one side consisted of clear, plexiglass on 3 sides and bright illumination of the area with no lid, identified as the “light” chamber. The other side consisted of four black plexiglass walls with a black lid, identified as the “dark” chamber. A camera was placed above the light-dark box to capture video footage for scoring. To begin the test, experimental mice were placed in the dark chamber against a corner and lid was immediately placed. Time spent in the light and dark chambers was recorded over the course of the 10-minute trial. Behavioral outcomes were automatically scored using Ethovision 18.0 and validated by a blinded scorer.

#### Tissue Collections

To collect brain tissue for further analysis, animals were sacrificed between PND46-PND50 via carbon dioxide inhalation for 5 minutes. The euthanasia was confirmed with a toe-pinch to ensure a lack of response to painful stimuli. Animals were weighed, and subsequently transcardially perfused with ice-cold saline for 4 minutes, followed by 3 minutes of 4% paraformaldehyde (PFA). Full brains were extracted and placed into glass vials with 5mL PFA. Brains were post-fixed in 4% PFA for 48hrs at 4°C, followed by cryoprotection in 30% sucrose with sodium azide for at least 48hrs at 4°C. Brains were then flash frozen using 2-methylbutane in dry ice and stored at −80°C until post-processed.

#### Cryosectioning and Immunohistochemistry

Brains were cryosectioned on a Precisionary CF-6100 Cryostat at a temperature of −20C, and a thickness of 30um per section. Brains sections in cryoprotectant were rinsed with PBS, mounted onto glass gelatin-subbed microscope slides and baked for 48-72 hrs at 37°C to encourage adherence. Tissue sections of interest containing the PVN were then stained. Tissue was first rinsed 3 x 5min with 1xPBS, followed by permeabilization with 1% triton-100x in PBS for 15 min at room temperature. Tissue was rinsed 3 x 5min with 1xPBS, followed by epitope retrieval via incubation of samples in 10mM citric acid (pH 9.0) at 70°C for 30 min. Tissue was removed from 70°C oven and left at room temperature to cool for 10 minutes, followed by 3 x 10min rinses with 1xPBS. Background fluorescence was quenched using 1mg/mL sodium tetraborate in 0.1M PB for 60 min at room temperature, followed by 6 x 5min rinses with 1xPBS. Tissue was then blocked with 10% normal goat serum (NGS; Vector Laboratories S-1000-20) in 1xPBS for 60 min at room temperature. Samples were subsequently incubated for 48 hours at 4°C in a solution consisting of 5% NGS, 0.3% Tween-20 in 1xPBS with primary antibodies. Primary antibodies used included: guinea pig anti-Iba1 (Synaptic Systems, 1:500), guinea pig anti-Tmem119 (Synaptic Systems, 1:500), mouse anti-vGlut2 (Synaptic Systems, 1:500), and rabbit anti-PSD95 (Life Technologies, 1:250). After 48 hours, samples were rinsed with 6x 10min with 1xPBS, followed by a 4hr incubation at room temperature in a solution consisting of 5% NGS, 0.3% Tween-20 in 1xPBS with secondary antibodies. The secondary antibodies used included: goat anti-guinea pig Alexa Fluor 568 (ThermoFisher Scientific, 1:200), goat anti-mouse Alexa Fluor 488 (ThermoFisher Scientific, 1:200), and goat anti-rabbit Alexa Fluor 647 (ThermoFisher Scientific, 1:200). Tissue was then rinsed with 6 x 10min with 1xPBS, followed by a 15 min incubation in in 1xPBS containing NucBlue Live cell stain (ThermoFisher Scientific; 1drop per 1000uL PBS) at room temperature. Samples were finally rinsed with a final round of 3 x 10min 1xPBS washes and subsequently cover slipped with VECTASHIELD Antifade Mounting Medium Plus, sealed with nail polish and placed in a dark container to seal overnight.

#### Confocal Microscopy and analysis for synaptic quantification and microglial morphology

To quantify excitatory synapse number and microglial engulfment of synaptic material, confocal microscopy images of the microglial markers Iba1 and Tmem119, the presynaptic excitatory marker vGlut2 and the postsynaptic marker PSD95 were acquired by a blinded observer using an Andor BC43 confocal microscope. Each image consisted of a z-stack of XY dimension 208.08 µm x 208.08 µm (2040 x 2040 pixels, 16 bit) using a 60x objective with an interval of 0.1 μm. Bitplane Imaris 10.0 volumetric reconstruction software was used to quantify excitatory synapse number and microglial engulfment of synaptic material. Specifically, Imaris’ Surface tool was used to reconstruct microglia (Iba1+/Tmem119+), vGlut2 puncta, and PSD95 puncta. Using Imaris’ object-object statistics function, excitatory synapses were identified as co-localized (≤ 0.0 μm distance) vGlut2 and PSD95 surfaces outside of reconstructed microglial or DAPI surfaces. Synaptic material was defined to be engulfed by microglia when the surface-to-surface distance between a vGlut2 or PSD95 puncta and the reconstructed microglial surface was < 1.0 x 10^-35^ μm. This threshold was selected to capture puncta located within the interior of microglial surfaces while accounting for the numerical precision limits of Imaris surface reconstruction, in which puncta that abut the inner boundary of a microglial surface are assigned a non-zero distance value rather than zero.

To quantify microglial morphology, 40x magnification Z-stacks of XY dimensions 314.16 µm x 314.16 µm (2040 x 2040 pixels, 16 bit) were taken every 0.3 μm on an Andor BC43 confocal microscope. Using FIJI, open microscopy environment tiffs were converted to 8-bit images. A blinded observer then used a MATLAB-based, semi-automated program to quantify microglial ramification index (CellSelect-3DMorph) (Taborda-Bejarano et al., 2025).

#### Statistical Analysis

All statistics and data visualization were performed using GraphPad Prism Version 11. A maximum of 2 pups per sex per litter were used to account for potential litter effects. Statistical tests performed for each experiment are reported in figure legends and included in Table 1. Individual points on graphs represent individual biological samples. Statistical significance was determined at p < 0.05. Partial eta squares (η^2^_p_) effect size and Cohen’s *f* effect size were calculated from GraphPad Prism-generated sum of squares.

## RESULTS

### Offspring Neonatal Behavioral Assays

We generated a TLR4 conditional knockout (TLR4 cKO) mouse line by crossing TLR4^flox/flox^ mice with *Cx3cr1*-BAC-Cre-transgenic mice (Ceasrine et al., 2022)(Fig. 1A-B). To determine whether microglial TLR4 signaling influences early communicative behavior, we recorded maternal separation-induced ultrasonic vocalizations (USVs) at postnatal day 8 (PND8). 2-way ANOVAs (genotype x sex) revealed a consistent main effect of genotype in both male and female offspring (Fig. 1C-1J and Supplemental Fig. 1). TLR4 cKO pups produced significantly fewer calls compared to Cre-negative TLR4^flox/flox^ littermate controls across nearly all quantified call subtypes, including step-down, step-up, frequency-modulated, chevron, complex, and multi-step. However, no significant impact of genotype or offspring sex was found for average call duration, average call pitch, or average call bandwidth (Supplemental Fig. 1). The breadth of this reduction across structurally distinct call types suggests that TLR4 cKO affects overall vocal output rather than selectively disrupting a specific acoustic feature of communicative behaviors.

**Figure 1.**
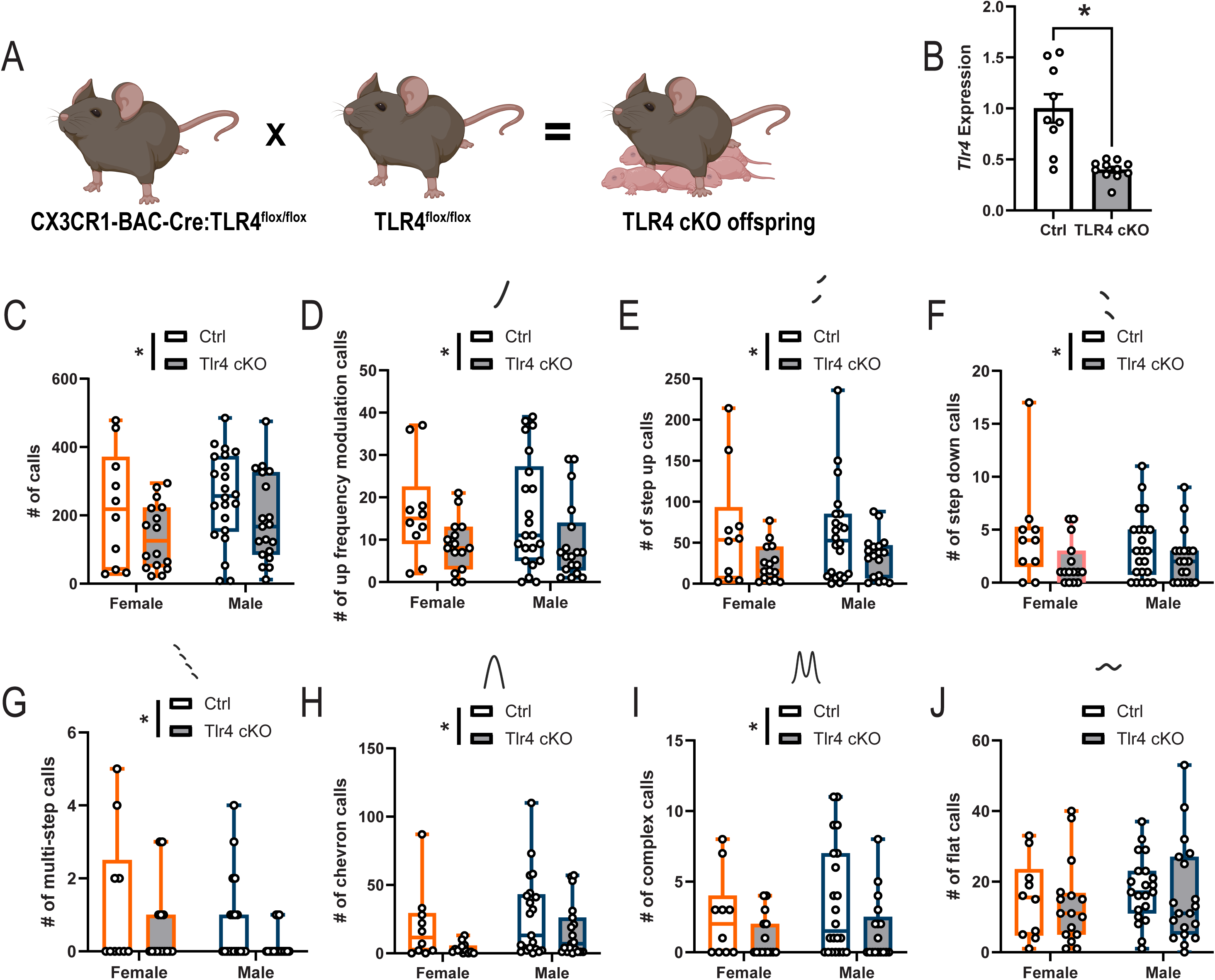
Deletion of TLR4 within CX3CR1-expressing cells alters neonatal communicative behaviors. **A.** Pictorial representation of breeding schema to generate TLR4 conditional knockout (TLR4 cKO) mice. **B.** Microglia (CD11b+ cells) were isolated from the forebrain of TLR4^flox/flox^:Cre^-^ (Ctrl) and TLR4^flox/flox^:Cre^+^ (TLR4 cKO) mice. qPCR for *Tlr4* relative to 18s confirmed conditional knockout of *Tlr4* expression in brain CX3CR1-expressing cells (p < 0.05, *d* = 1.44). Differences between groups were assessed by two-tailed t-test, effect size was assessed by Cohen’s *d*. Bar graphs display mean ± standard error of the mean. * p < 0.05. N = 9-11 per group. **C-J.** Maternal separation-induced ultrasonic vocalizations (USVs) were assessed on postnatal day 8 in Ctrl and TLR4 cKO mice and analyzed using VocalMat software. VocalMat-produced call depictions are represented above each respective call type. **(C)** Number of total calls, **(D)** number of up frequency modulation calls, **(E)** number of step up calls, **(F)** number of step down calls, **(G)** number of multi-step calls, **(H)** number of chevron calls, **(I)** number of complex calls, and **(J)** number of flat calls were quantified. Differences between groups were assessed by 2-way ANOVA (sex x genotype). Bar graphs display mean ± standard error of the mean. Significant main effects of genotype are depicted by * p < 0.05. N = 9-22 per group. Figure was created in part using Biorender.

### Offspring Adolescent Behavioral Assays

Litters were weaned on PND24 after which adolescent behavioral testing batteries began on PND30. Importantly, TLR4 cKO did not significantly alter offspring weight across early development (Supplemental Fig. 2). We first assessed social preference using a modified Crawley’s 3-chambered arena task (Fig. 2A) in which the preference for a test mouse to investigate either a novel age- and sex-matched conspecific or a novel object was determined (a rubber duck was used as a standardized non-social stimulus). Conditional knockout of TLR4 within CX3CR1-expressing cells had no impact on social motivation in male or female offspring (Fig. 2B-C), although we did observe a main effect of sex on % sociability, with males displaying greater sociability. We also assessed anxiety-like behaviors using the elevated zero maze and light/dark box assays. No significant differences in anxiety-like behavior were observed in either test (Fig. 2D-F)

**Figure 2.**
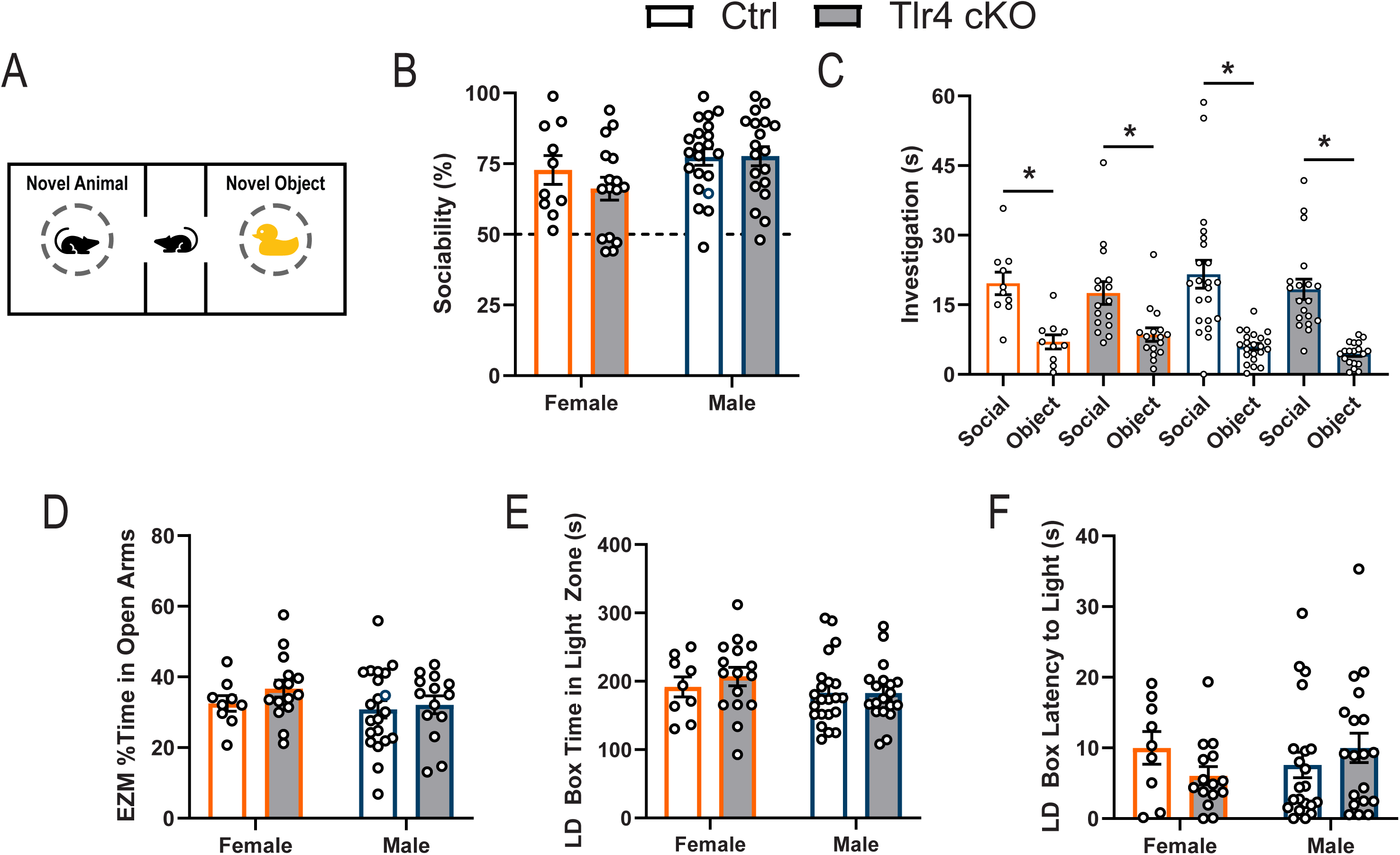
Adolescent social and anxiety-like behaviors do not require microglial TLR4. **A.** Female and male mice were placed in a 3-chambered arena and given the choice to interact with either a novel age- and sex-matched conspecific mouse or an inanimate object for 5 min on PND30. **B.** Time spent investigating the novel animal compared to the novel object (% sociability) was assessed Differences across groups were assessed by 2-way ANOVA (sex x genotype). Bar graphs display mean ± standard error of the mean. N = 10-21 per group. **C.** Social investigation time and novel object investigation time was compared within female and male Ctrl and Tlr4 cKO mice. Differences across groups were assessed by 1-way ANOVA followed by Bonferroni’s post-hoc. * p < 0.05. N = 10-22 per group. **D.** Mice were placed in an elevated zero maze for 5 min and the percentage of time spent in the open arms was assessed. Differences across groups were assessed by 2-way ANOVA (sex x genotype). Bar graphs display mean ± standard error of the mean. N = 10-21 per group. **E-F.** Mice were placed in a light dark box for 10 min and **(E)** the time spent in the light zone and **(F)** the latency to enter the light zone were assessed. Differences across groups were assessed by 2-way ANOVA (sex x genotype). Bar graphs display mean ± standard error of the mean. N = 10-22 per group.

### Microglial Morphology and Phagocytosis

To assess whether the constitutive downregulation of TLR4 in CX3CR1-expressing cells impacted microglial morphology, we used a MATLAB-based, semi-automated program (CellSelect-3DMorph) (Taborda-Bejarano et al., 2025) to quantify the microglial ramification index of microglia in the PVN (Fig. 3A). Conditional knockout of TLR4 in CX3Cr1-expressing cells robustly increased both the ramification index (Fig. 3B) as well as the cell volume (Fig. 3C) of Iba1+/Tmem119+ microglia in the PVN of both male and female offspring. This increase in ramification, reflecting a more elaborate, branched morphology, suggests that TLR4 signaling may normally constrain microglial process extension under homeostatic conditions, and that its loss shifts microglia towards a morphologically distinct state.

**Figure 3.**
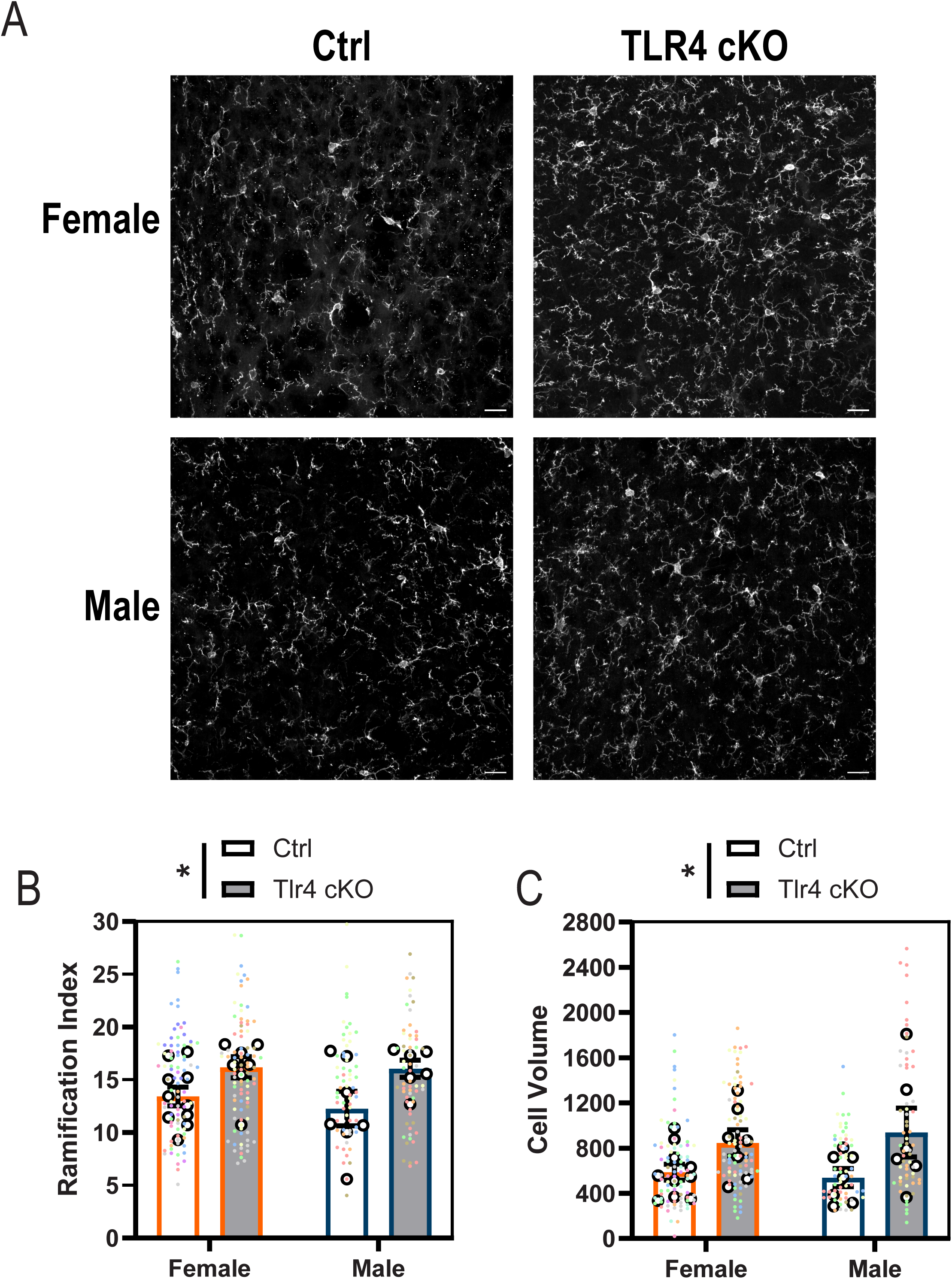
Ablation of TLR4 in CX3CR1 cells alters microglial morphology. **A.** Representative 40x images of microglia (Iba1+Tmem119+ cells) in the paraventricular nucleus (PVN) of the hypothalamus. Representative images for female and male Ctrl and TLR4 cKO PVN are shown. Scale bar = 20 μm. **B-C. (B)** Ramification index and **(C)** cell volume of microglia within the PVN was assessed. Bar graphs display mean ± standard error of the mean. Super plots where biological means are depicted by open white circles and smaller dots where each color is indicative of individual microglia from that animal. Differences between groups were assessed by 2-way ANOVA (sex x genotype). Significant main effects of genotype are depicted by * p < 0.05. N = 6-10 per group.

Given the robust morphological alterations observed in TLR4 cKO microglia, we next assessed whether the capacity of microglia to engulf synaptic material in the PVN was changed in the absence of TLR4. To do so, we used confocal microscopy and IMARIS volumetric reconstruction software to assess vGlut2+ synapses as well as engulfment of synaptic material in the PVN. Conditional knockout of TLR4 did not significantly alter the total number of excitatory synapses in the PVN (Fig. 4). However, there was a significant main effect of genotype on microglial engulfment of vGlut2+ presynaptic puncta, with a more robust decrease in engulfment observed in males than in females (Fig. 4). No significant alterations were observed in microglial engulfment of PSD95+ puncta.

**Figure 4.**
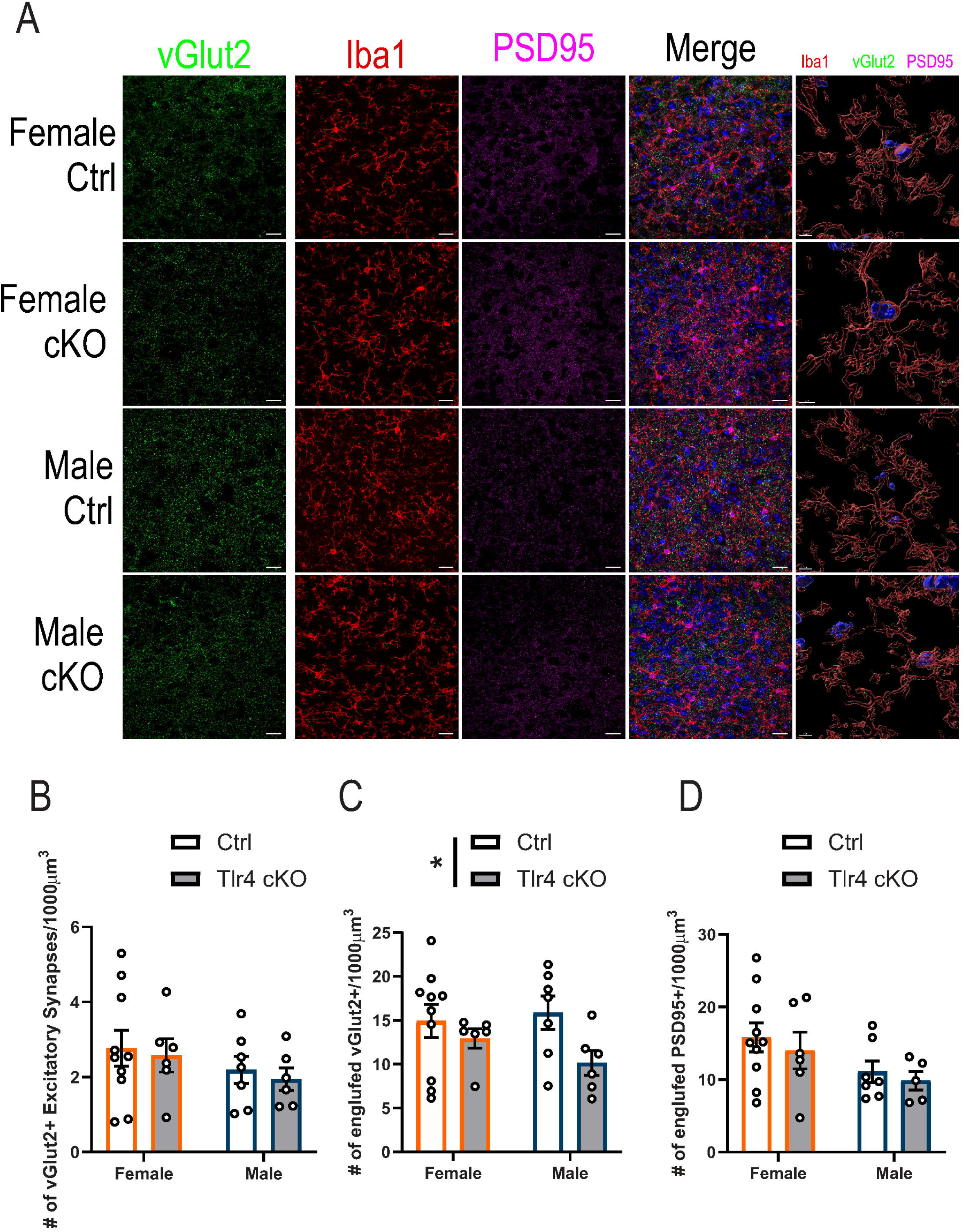
Microglial engulfment of presynaptic excitatory material is altered by TLR4 knockout. **A.** Representative 60x images of vGlut2, PSD95, and microglia (Iba1+Tmem119+ cells) in the paraventricular nucleus (PVN) of the hypothalamus. Representative images for female and male Ctrl and TLR4 cKO PVN are shown. An enlarged image of the IMARIS volumetric reconstruction demonstrating DAPI (blue), along with vGlut2 (green) and PSD95 (magenta) within microglia (red). Scale bar for 4 left columns = 20 μm. Scale bar for rightmost column = 5 μm. **B.** Number of vGlut2+/PSD95+ excitatory synapses in the PVN was quantified. **C.** The number of vGlut2+ puncta contained within Iba1+Tmem119+ microglial surfaces was quantified. **D.** The number of PSD95+ puncta contained within Iba1+Tmem119+ microglial surfaces was quantified. Differences across groups were assessed by 2-way ANOVA (sex x genotype). Bar graphs display mean ± standard error of the mean. Significant main effects of genotype are depicted by * p < 0.05. N = 6-10 per group.

Because early-life adversity has been shown to disrupt microglial engulfment in the PVN (Bolton et al., 2022), we assessed whether conditional knockout of TLR4 within CX3CR1-lineage cells would alter this response. To do so, we exposed TLR4 cKO and Cre-negative TLR4^flox/flox^ littermates to LBN from PND2-10 (Supplemental Fig. 2). Consistent with previous literature, maternal exposure to this LBN paradigm significantly altered maternal behavior towards offspring (Walker et al., 2017; Bolton et al., 2022), as LBN-exposed mothers spent significantly less time on the nest taking care of their litters, and significantly more time in non-nursing/grooming-related behaviors (Supplemental Fig. 2). Previous literature assessing the impact of LBN on microglial and synaptic outcomes has focused on earlier timepoints and specifically on CRH-expressing neurons in the PVN (Bolton et al., 2022; Garvin et al., 2025). Here, we assessed the impact of early life adversity induced by LBN caging and TLR4 cKO in males and females early in adulthood, following the completion of a behavioral battery. Notably, there is a robust effect of LBN on vGlut2+ excitatory synapse number that is ameliorated by TLR4 cKO in male mice (significant genotype x sex x caging interaction) (Supplemental Fig. 4). Microglial morphological analyses also revealed a significant interaction between genotype and treatment on ramification index (Supplemental Fig. 3).

## DISCUSSION

Here, we demonstrate that conditional knockout of TLR4 in CX3CR1-expressing myeloid cells is sufficient to alter microglial morphology, reduce microglial engulfment of presynaptic material in the PVN, and suppress early communicative behaviors under standard housing conditions, even in the absence of exogenous immune challenges or adversity. LBN studies suggest that TLR4 conditional knockout may also modify microglial responses to early life stress. Collectively, these findings suggest that the role of TLR4 within microglia extends beyond its canonical functions in pathogen detection, contributing to the regulation of cell morphology, phagocytic capacity, and neurodevelopmental outcomes during healthy brain development.

One of the most unexpected findings of this study was that reduced microglial engulfment of presynaptic material occurred without a corresponding reduction in excitatory synapse density within the PVN. Although conditional knockout of TLR4 increased microglial ramification and cell volume, these morphological alterations were accompanied by a decrease, rather than the expected increase, in engulfment of vGlut2 positive presynaptic punctate. This difference suggests that microglial morphology alone may not consistently predict phagocytic capacity during postnatal development. The absence of a detectable change in overall excitatory synapse number may indicate that compensatory pruning pathways operate in parallel to microglial TLR4-dependent mechanisms, or that alterations in synaptic turnover occurred earlier in development and were no longer detectable at the timepoint examined in this study. Together, these findings reveal an unexpected disentangling of microglial engulfment and excitatory synapse density, suggesting that reduced TLR4-dependent phagocytosis alone may be insufficient to alter mature synaptic architecture within the PVN.

The selective reduction in engulfment of vGlut2+ puncta, without a corresponding change in the engulfment of PSD95+ puncta, is consistent with previous studies demonstrating preferential complement-mediated elimination of presynaptic inputs during postnatal development (Stevens et al., 2007; Schafer et al., 2012). Given the known cross-talk between TLR4 activation and complement component expression (Pope et al., 2010; Song, 2012; Hajishengallis and Lambris, 2016; Yang et al., 2020), tonic microglial TLR4 activity may support complement-dependent pruning. However, it must be noted that we did not directly assess complement deposition in this study. Future studies examining C1q and C3 colocalization with vGlut2+ puncta in TLR4 cKO versus control PVN at early postnatal timepoints will be necessary to test this hypothesis and to determine whether the engulfment deficit we observed is complement-dependent or instead reflects TLR4-dependent regulation of alternative phagocytic pathways such as MerTK or TREM2 (Scott-Hewitt et al., 2020; Garvin et al., 2025).

That TLR4 cKO pups showed a reduction in maternal separation-induced USVs across structurally diverse call subtypes is consistent with a growing literature demonstrating that microglial depletion or dysfunction during the perinatal period alters both synaptic connectivity and behavioral outcomes, and with evidence that developmental TLR4 manipulation alters offspring social and communicative behavior (Zhan et al., 2014; Kopec et al., 2018; Xiao et al., 2021). Analyses were performed on the PVN because of its established role in stress-sensitive microglial pruning (Bolton et al., 2022; Garvin et al., 2025). However, neonatal vocalization depends upon distributed brainstem, hypothalamic, and forebrain circuits (Michael et al., 2020; Park et al., 2024; Chen et al., 2026) and therefore, the behavioral phenotype cannot be attributed solely to altered PVN circuitry. The restriction of behavioral impact to the neonatal period may reflect the well-documented capacity of postnatal circuits to compensate for early perturbations during sensitive periods.

While *Cx3cr1*-Cre animals are often discussed as microglia-specific tools, CX3CR1 is expressed within a variety of tissue resident myeloid cells, including intestinal lamina propria macrophages that mediate gut immune homeostasis and microbiota sensing. Germ-free mice lacking commensal microbiota exhibit global microglial defects, including altered cell proportions, an immature phenotype, and impaired innate immune responses (Erny et al., 2015). Germ-free studies have demonstrated that gut-derived microbiota can influence microglial immune function to prevent brain damage following viral infection, likely in a TLR4-dependent manner (Brown et al., 2019). Consistent with the broader principle that early microbial environments shape brain development, germ-free neonates exhibit altered microglial densities within the PVN, hippocampus, and somatosensory cortex (Castillo-Ruiz et al., 2023). Given this, our findings cannot fully distinguish a brain-intrinsic contribution of microglial TLR4 from a peripheral contribution arising from TLR4 knockout in intestinal CX3CR1-expressing myeloid cells. Disentangling these possibilities will require truly microglia-specific approaches.

Taken together, the present findings advance a revised understanding of TLR4 function in the postnatal CNS. Rather than serving exclusively as a sentinel for infection or tissue injury, TLR4 within CX3CR1-expressing myeloid cells contributes to microglial morphology, presynaptic engulfment, and neonatal communicative behavior under basal conditions. These findings support a broader conceptual framework in which innate immune receptors are not simply activated during pathological states but also participate in the developmental programs which shape circuit maturation. Therefore, disruption of this innate immune signaling, even in the absence of an overt inflammatory event, may alter the trajectory of normal brain development through mechanisms extending beyond classical immune defense.

### Limitations

While we demonstrate TLR4 downregulation in microglia from cKO animals, we cannot exclude residual TLR4 expression in a subset of CX3CR1-expressing cells or functional compensation through other pattern recognition receptors (i.e. TLR2), or alterations in CX3CR1 cells outside of the CNS. Second, the *Cx3cr1*-BAC-Cre driver targets not only microglia but also peripheral monocyte-derived cells that may transiently enter the brain during postnatal development; the contribution of peripheral myeloid TLR4 deletion to the observed phenotypes cannot be entirely excluded, though recent characterization of this line suggests minimal off-target recombination (Mroue-Ruiz et al., 2026). Third, the mechanistic link between reduced TLR4 signaling, altered microglial morphology, impaired presynaptic engulfment, and suppressed USVs remains correlative; whether these phenotypes are causally connected, and through which downstream signaling intermediaries may be connected, remains unresolved.

## Supporting information

Statistics Table

