## Supplemental Materials and Figures for "*Cx3cr1*-BAC-CRE-mediated knockout of toll-like receptor 4 alters mouse communicative behaviors, microglial morphology, and engulfment of synaptic material"

#### SUPPLEMENTAL METHODS

##### Limited Bedding and Nesting Paradigm

To introduce the early life stress variable of Limited Bedding and Nesting (LBN), dams were randomly assigned either LBN or control housing conditions (Supplemental Fig. 2). Control housing consisted of a standard polypropylene cage with cobb bedding, a cotton nestlet (1in x 1in) in place of CareFresh, and with an igloo for enrichment. LBN housing consisted of a standard polypropylene cage with cobb bedding atop of which was placed a wire mesh (McNichols expanded metal Item No 4700313244) separating mice from access to cobb bedding, as well as half of a cotton nestlet (1in x 1in triangle), and with an igloo. The wire mesh was laid flat in the bottom of the polypropylene cage to separate mice from access to cobb bedding, while maintaining maximal cleanliness secondary to urine and feces. Dams and their pups were placed into their randomly assigned cage condition on the morning of PND2 and remained in this cage until the morning of PND10 (Bolton et al., 2022) at which point all experimental cages were exchanged for identical fresh polypropylene cages with cobb bedding, a cotton nestlet (1in x 1in) in place of CareFresh, and with an igloo for enrichment. This concluded the LBN early life stress paradigm, and all animals remained in standard cages with nestlets until offspring reached weaning age, at which point all animals were placed in standard polypropylene cages (27 cm x 16 cm x 15.5 cm) with cobb bedding and CareFresh.

#### SUPPLEMENTAL FIGURE LEGENDS

**Supplemental Figure 1. A-G.** Maternal separation-induced ultrasonic vocalizations (USVs) were assessed on postnatal day 8 in Ctrl and TLR4 cKO mice and analyzed using VocalMat software. **(A)** Number of down frequency modulation calls, **(C)** number of reverse chevron calls, **(C)** number of two step calls, **(D)** number of short calls, **(E)** average call duration, **(F)** average call pitch, and **(G)** average call bandwidth were quantified. Differences between groups were assessed by 2-way ANOVA (sex x genotype). Bar graphs display mean ± standard error of the mean. Significant main effects of genotype are depicted by * $p < 0.05$. N = 9-22 per group.

**Supplemental Figure 2. A-C.** Pictorial representation of early-life adversity paradigm. To model early-life resource deprivation, we placed dams and litters into either **(A)** standard (Ctrl) housing with cobb bedding and a cotton nestlet or **(B)** into a limited bedding and nesting (LBN) environment with wire mesh flooring, limited bedding material, and half of a cotton nestlet from postnatal days 2-9. **(C)** Maternal care observations were performed on postnatal days 3, 6, and 9. Ultrasonic vocalizations (USVs) were recorded on postnatal day 8. Litters were returned to standard housing environments on postnatal day 10. All litters were weaned on postnatal day 24. Adolescent behavioral testing paradigm began on postnatal day 30. Differences across groups. **D.** Maternal care was assessed at postnatal days 3, 6, and 9 in the home cage. 30 observations were made over 30 minutes, and the number of times the dam was off nest during this time period was reported. **E.** Offspring weights were assessed on postnatal days 8, 12, 24, and 30. Differences across groups were assessed by a mixedeffects REML model (based on a 3-way repeated measures ANOVA) followed by Bonferroni's post-hoc. * $p < 0.05$. N = 10-22 per group.

**Supplemental Figure 3. Ablation of TLR4 in CX3CR1 cells alters microglial morphology. A.** Representative 40x images of microglia (Iba1+Tmem119+ cells) in the paraventricular nucleus (PVN) of the hypothalamus. Representative images for female and male Ctrl and TLR4 cKO PVN are shown. **B-C. (B)** Ramification index and **(C)** cell volume of microglia within the PVN was assessed. Bar graphs display mean ± standard error of the mean. Super plots where biological means are depicted by open white circles and smaller dots where each color is indicative of individual microglia from that animal. Differences between groups were assessed by 3-way ANOVA (offspring sex x genotype x housing condition) followed by Bonferroni's post-hoc. N = 6-10 per group. Scale bar = 20 μm.

**Supplemental Figure 4. Impact of early life adversity of excitatory synapses and microglial synaptic engulfment in mice lacking myeloid cell TLR4. A.** Representative 60x images of vGlut2, PSD95, and microglia (Iba1+Tmem119+ cells) in the paraventricular nucleus (PVN) of the hypothalamus. Representative images for female and male Ctrl and TLR4 cKO PVN are shown. **B.** Number of vGlut2+/PSD95+ excitatory synapses in the PVN was quantified. **C.** The number of vGlut2+ puncta contained within Iba1+Tmem119+ microglial surfaces was quantified. **D.** The number of PSD95+ puncta contained within Iba1+Tmem119+ microglial surfaces was quantified. Differences across groups were assessed by 3-way ANOVA (offspring sex x genotype x housing condition) followed by Bonferroni's post-hoc. Bar graphs display mean ± standard error of the mean. * $p < 0.05$. N = 6-10 per group. Scale bar for 4 left columns = 20 μm. Scale bar for rightmost column = 5 μm.

### Supplemental Figure 1.

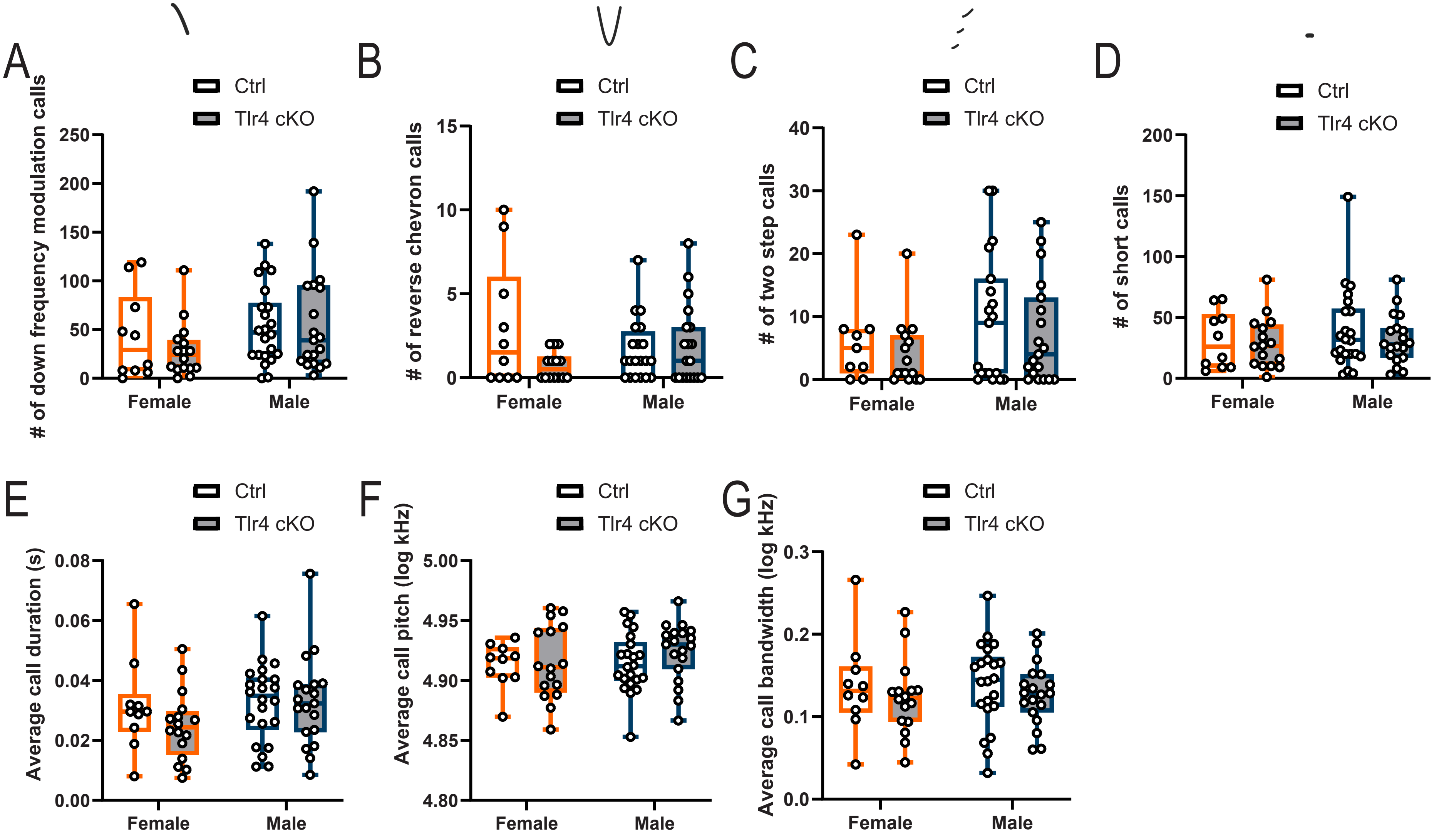

**Supplemental Figure 2.**

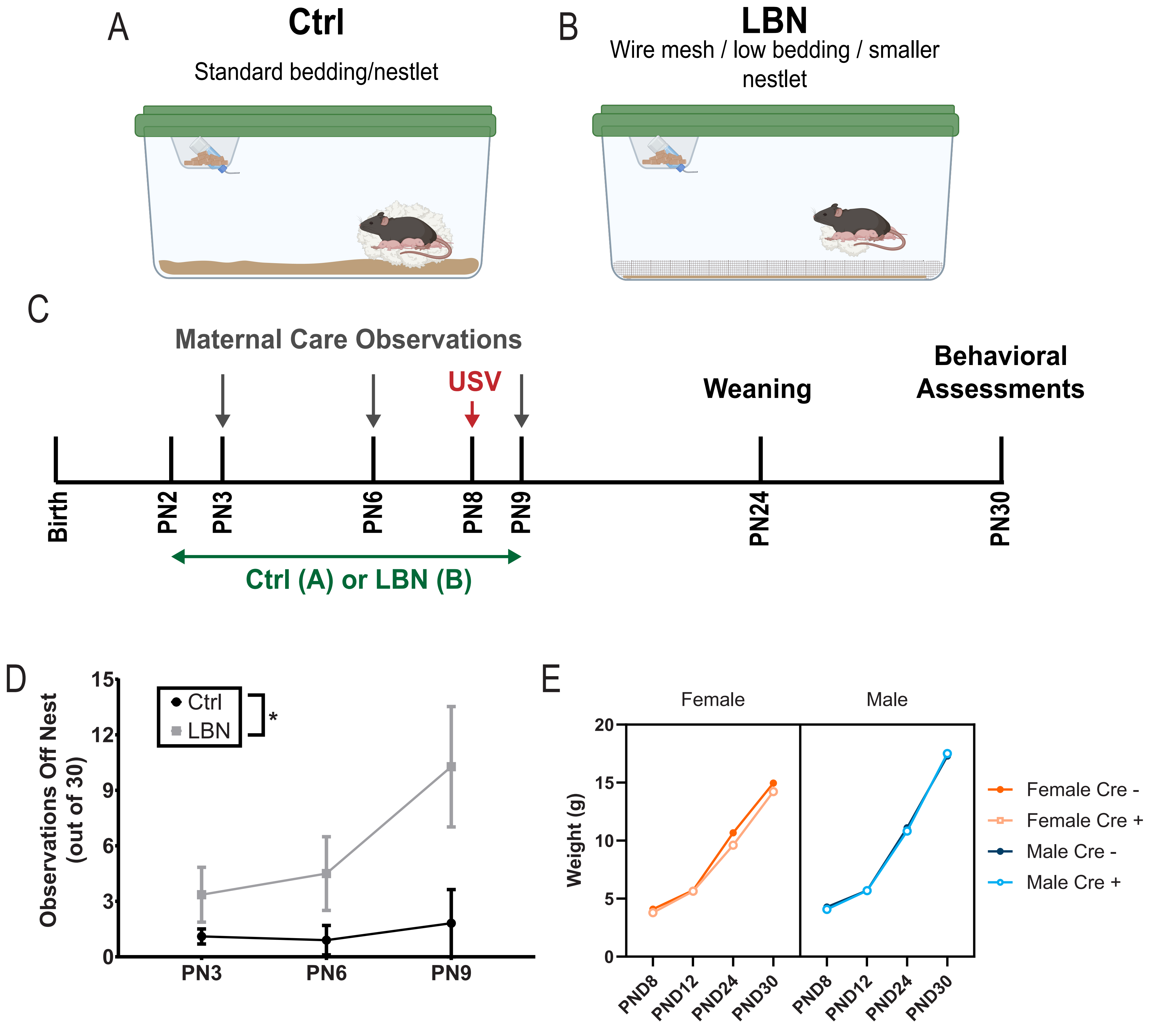

**Supplemental Figure 3.**

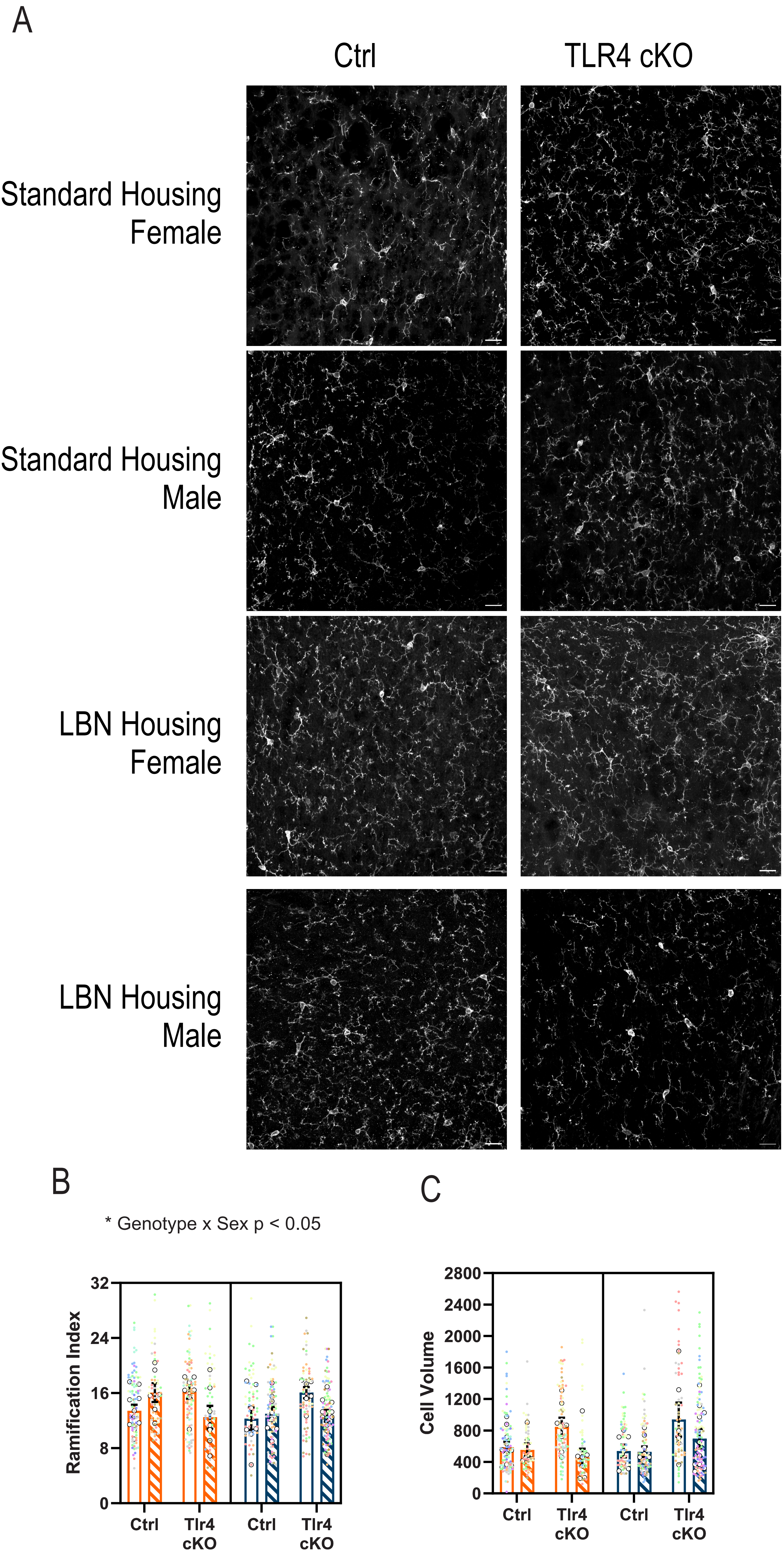

**Supplemental Figure 4.**

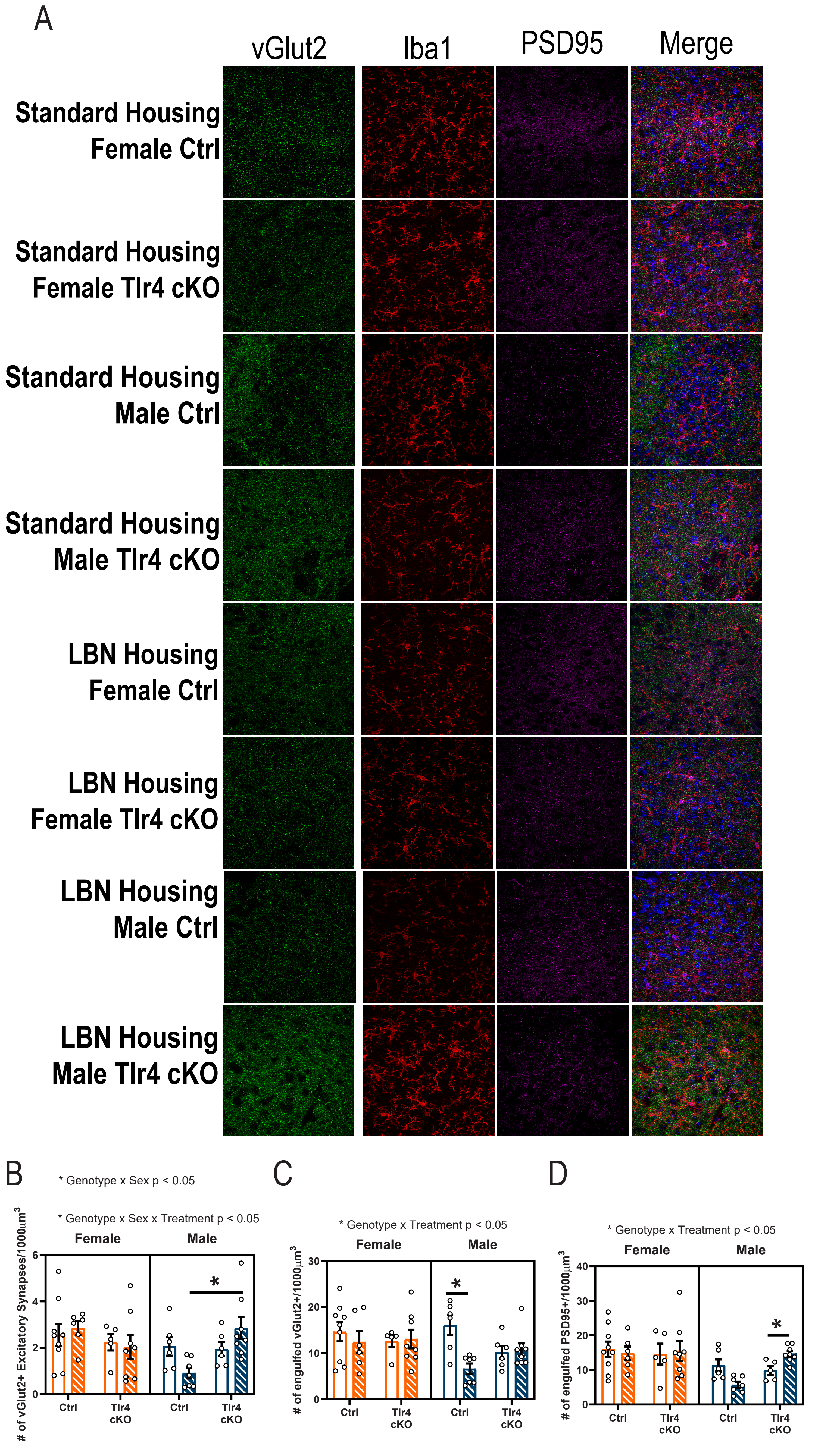
